# Perinatal circadian desynchronization disrupts sleep and prefrontal cortex function in adult offspring

**DOI:** 10.1101/2025.01.21.634103

**Authors:** Brandon L. Roberts, Jiexin Wang, Haifa Chargui, Nathan C. Cupertino, Walker Sorensen, Ilia N. Karatsoreos

**Author notes:** **Correspondence should be sent to:** Brandon L. Roberts, Ph.D., Department of Zoology & Physiology, University of Wyoming, 1000 E. University Ave, Biological Sciences, Room 316A, Laramie, WY 83071.

## Abstract

Sleep and circadian (daily) rhythms impact nearly all aspects of physiology and are critical for optimal organismal function. Disruption of the clock can lead to significant metabolic disorders, neuropsychiatric illness, and cognitive dysfunction. Our lab has previously shown that environmental circadian desynchronization (ECD) in adults alters the anatomical structure and neurophysiological function of prefrontal cortex (PFC) neurons, PFC mediated behaviors, as well as sleep quality. As the PFC undergoes significant development in utero and early life, and maternal disturbances during this period can have significant long-term ramifications, we hypothesized that disrupting the circadian environment of dams during the perinatal period would alter sleep and PFC function in adult offspring. Using a mouse model of ECD we investigated how perinatal ECD (pECD) modulates sleep quality in adult offspring. We also determined how pECD impacts PFC neural function in adult offspring using *ex vivo* patch-clamp electrophysiology, exploring how pECD alters synaptic function and action potential dynamics. We found that male pECD mice trended toward increased total sleep during the inactive (light) period with shorter sleep bouts during the active (dark) period. pECD did not change sleep behavior in female mice. Independent of time of day, pECD altered post-synaptic dynamics of excitatory neurotransmitter release onto plPFC pyramidal neurons. There was also a loss of time-of-day effects on cell endogenous properties in male pECD mice. Thus, pECD clearly alters sleep behavior and PFC function in male mice. However, female mice appear protected against the effects of pECD in these measures. Together, these experiments form the foundation for future studies to understand the lifelong neurobehavioral impact of pECD.

## Introduction

The circadian (daily) timing system is critical for optimal function of physiology and behavior. How perturbation of circadian timing during critical periods of development can impact later function is therefore an important question. It is well established that maternal environment heavily impacts offspring physiology, neurodevelopment, and causes epigenetic changes in offspring that can be passed on to future generations ^1–3^. Our group has demonstrated that environmental circadian desynchronization (ECD), induced by changing the light cycled to 10h light/10h dark (T20), negatively impacts metabolic function, emotionality, and cognitive flexibility^4^. However, it is unknown how exposure to ECD during the perinatal period impacts the neural circuits underlying emotional and cognitive behaviors. Identifying how the perinatal circadian environment impacts offspring later in life is essential if we are to understand the developmental components of physiological and psychological pathologies in adulthood.

The prefrontal cortex (PFC) is important in the regulation of many different behaviors, from executive function and cognitive flexibility to mood and emotional regulation^4,5^. Our lab has documented that prelimbic layer 2/3 PFC pyramidal neurons show remarkable changes in fundamental neurophysiological properties over the day, and that these effects differ between males and females^6^. Importantly, we have also shown that ECD in adulthood leads to cognitive rigidity, altered PFC morphology, and physiological perturbations in neural activity ^4,6^. Given the important role daily rhythms seem to play in normal PFC function, and that the PFC undergoes significant development *in uter*o and early life^7–9^, a current gap in knowledge is understanding how perinatal ECD (pECD) impacts the function of PFC neurons of offspring later in life.

Sleep homeostasis is a critical property of maintaining proper health^10^. In addition to ECD altering PFC function and associated behaviors, ECD also results in poor sleep quality, a potential contributor to our previous findings that ECD alters PFC function. However, it is unknown how perturbations in circadian environment during the perinatal period may influence sleep behavior and quality later in life. Here we use a novel mouse model of perinatal ECD (pECD) by changing the light cycle to T20 during the perinatal period. Offspring are weaned into a normal 24h light/dark cycle (T24) and left uninterrupted until adulthood, where they are evaluated for changes in sleep behavior and time-of-day dependent changes in neural function.

Here, we combine *in vivo* sleep monitoring with *ex vivo* patch-clamp electrophysiology, forming the foundation for future studies to understand the lifelong neurobehavioral impact of pECD. The primary outcomes of this study are that early life perturbations of daily rhythms have long-term sex-specific consequences for body weight, sleep, and neural function of PFC pyramidal neurons.

## Results

### Maternal environmental circadian desynchronization (pECD) decreases offspring body weight

Perturbations in circadian rhythms have consequences for metabolic function. However, the long-term impact of circadian desynchronization during the perinatal period in offspring was unknown. To determine how pECD impacts offspring physiology and behavior we used an pECD model where dams are placed into a 10:10 LD (T20) cycle two weeks prior to breeding to disrupt their daily rhythms ^4^. Dams are then bred in ECD conditions and pups remain with their mother until weaning. At weaning (P21), pups are placed into a standard 12:12 LD (T24) cycle and housed normally until adulthood (P47; **Fig. 1A**). We recorded the total number of pups born in CTR and pECD conditions and found no difference in litter size or impact of multiparity litter number on litter size (**Fig. 1B**). Further, we measured body weights of male and female pups every 4-6 days starting at time of weaning on postnatal day (P21) into adulthood (P47). Here we show that pECD decreases body weight in male and female mice, an effect that lasts into early adulthood (**Fig.1C-D**). Interestingly, while there was a main effect of the pECD treatment in male mice, posthoc comparisons revealed that the effect was only statistically significant at P47 (**Fig. 1C**). Female mice displayed a stronger effect early in development, but their body weight rebounded early in adulthood, returning to CTR weights by P42 (**Fig.1D**). This data suggests that there are long-term physiological consequences associated with exposure to circadian desynchronization early in life.

**Figure 1.**
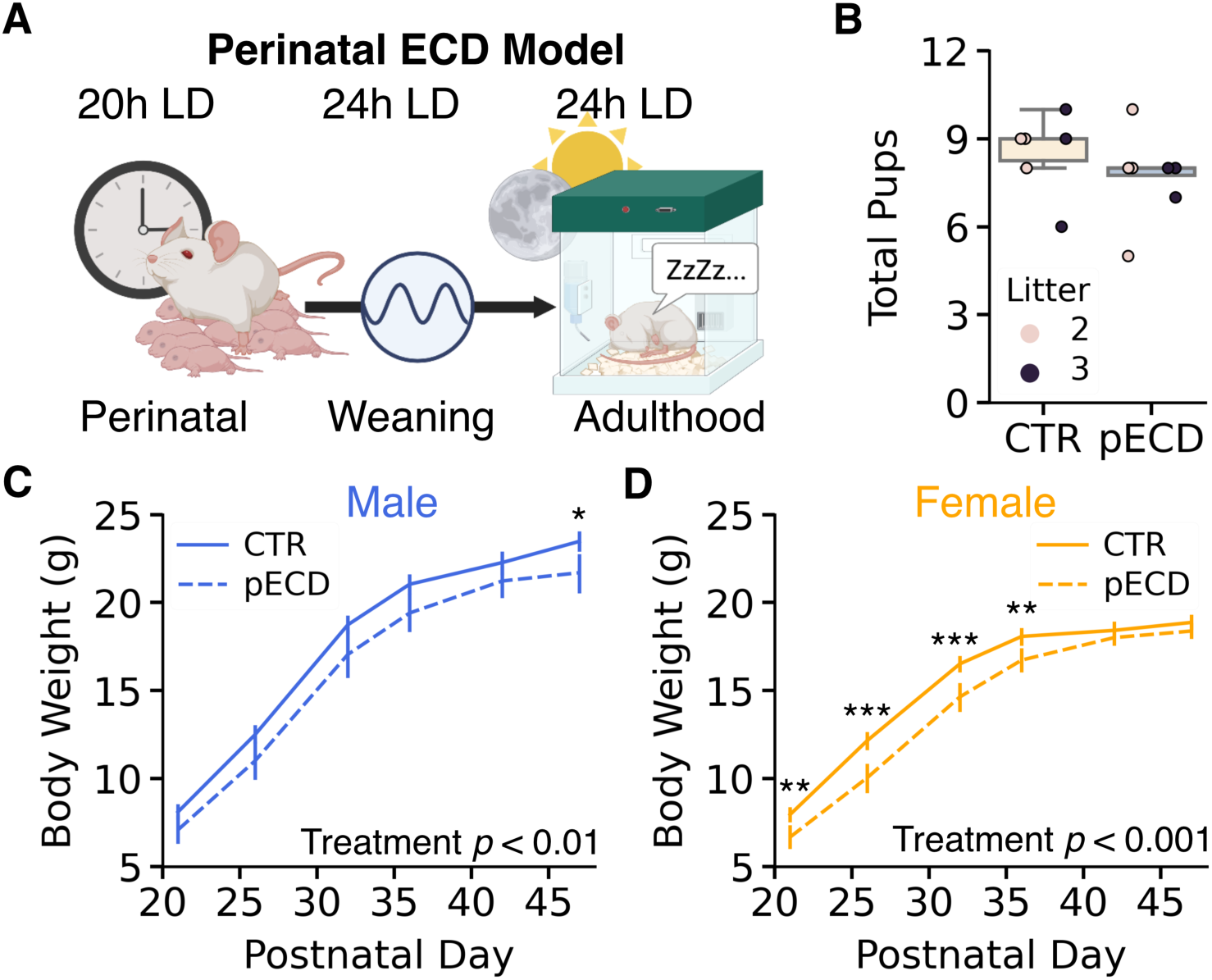
pECD does not affect litter size but does reduce body weight. ***A***, Diagram of perinatal circadian desynchronization (pECD) model. ***B,*** Total pups per litter born to control (CTR) and pECD dams (Treatment: F_(1, 10)_ = 0.90, *p* = 0.37; Litter number: *F_(1,10)_ = 0.07, p = 0.79*), ***C,*** Mean body weight of male *(Treatment: F_(1,32)_ = 9.18, p = 0.005; Age: F_(5,160)_= 2142.09, p < 0.001)* and (***D***) female *(Treatment: F_(1,41)_ = 18.97, p < 0.0001; Age: F_(5,205)_= 1593.81, p < 0.0001)* offspring from CTR and pECD dams between weaning at postnatal day (P) 21 and P47 (adulthood). N = 5-6 litters/group*. Two-way or mixed ANOVA for main effects (between treatment and/or litter, within age); ***p <0.001, **p <0.01, *p < 0.05*

### pECD does not impact total sleep in male or female mice

To test how pECD impacts sleep in adult pECD offspring, adult mice were placed in a non-invasive sleep monitoring system for 5 consecutive days^15,16^. The first 48h was used as an acclimation period and sleep measurements were analyzed using data from the last 72h in the system (**Fig.2A-D**). These data were combined to calculate average 24h (daily) sleep behavior. In male offspring, pECD did not impact average length of sleep bouts (**Fig. 2E**) or total sleep (**Fig. 2F**). We did not observe a difference in general measures of 24h sleep behavior in female mice, such as sleep bout length (**Fig. 2G**) or total sleep over the 24h period (**Fig. 2H**). Together this suggests that over the 24h period, pECD does not impact total sleep or bout length in male or female mice.

**Figure 2.**
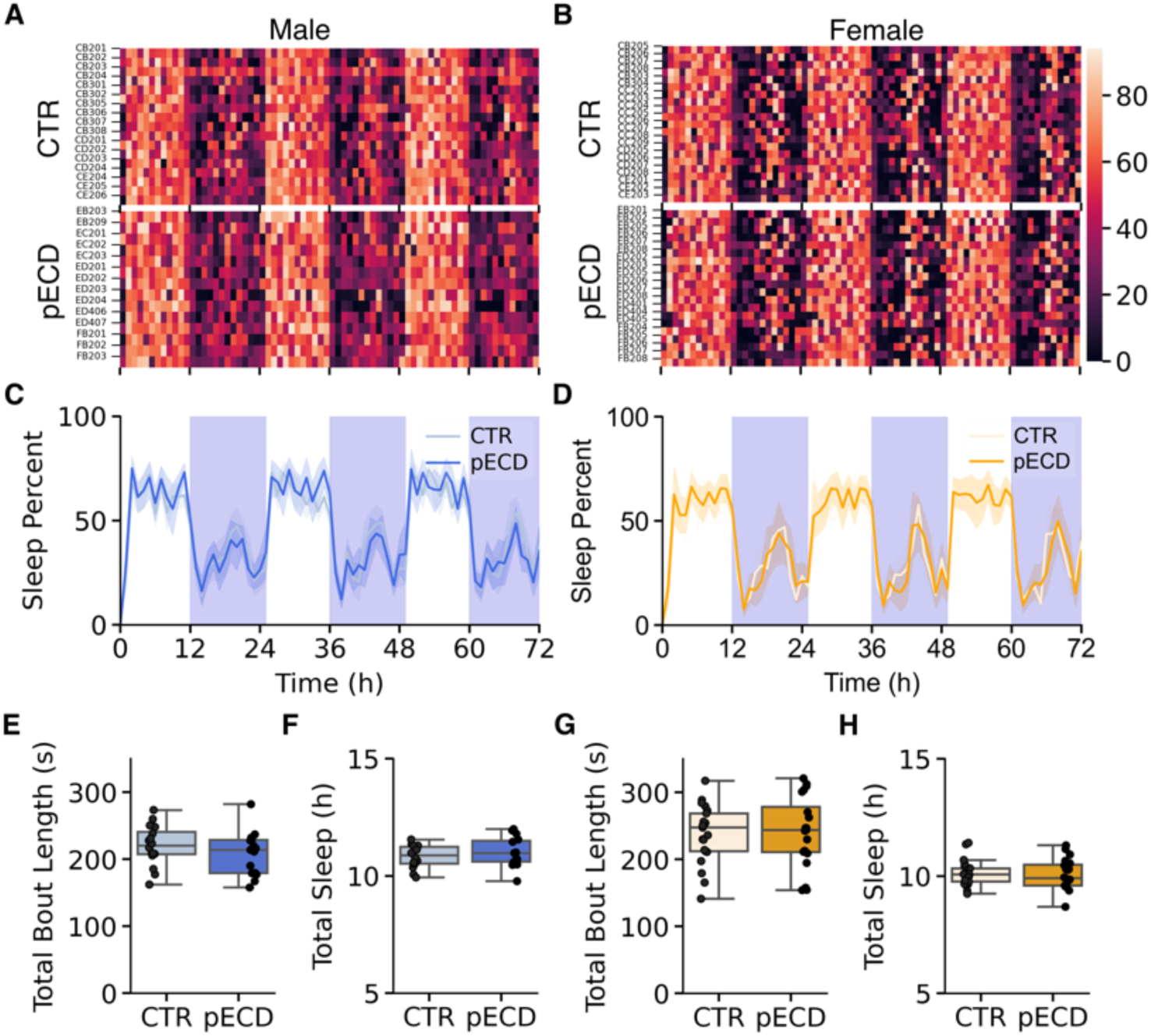
pECD does not affect coarse measures of sleep in either sex. ***A,*** Heat map of sleep behavior in individual CTR and pECD male and (***B***) female mice plotted in 1h bins sharing an x-axis with (***C,D***). ***C***, Continuous 72h sleep recordings averaged from all male and ***D*,** female CTR and pECD mice. (***E,F***) Boxplot and individual values averaged over 72h for daily sleep bout length (s) and total sleep (h) in male (Total Bout Length: t(29) = 1.09, *p* = 0.29; Total Sleep: t(29) = 0.86, *p* = 0.40) and (***G,H***) female (Total Bout Length: t(39) = 0.26, *p* = 0.79; Total Sleep: t(39) = 0.22, *p* = 0.83) mice (*n* = 14-21 mice/group). *Student t-test. Boxplots represent median, inner, and outer quartile range*.

### Male pECD offspring have fragmented sleep during the dark and sleep more during the light

Mice typically sleep in short bouts on the order of seconds and minutes, with more sleep during the light (inactive) phase and less sleep during the dark (active) phase. Although there was no difference in mean bout length or total sleep over the 24h period (**Fig. 2**), an important component of quality sleep is how it is distributed between the active and inactive period, and if this sleep is fragmented into short sleep bouts or consolidated into longer sleep bouts^16,17^. To test how pECD may impact sleep at the granular level in adult offspring we measured how their sleep during the light and dark period was distributed across eight separate bin sizes, ranging from 16 to 2048 seconds (**Fig. 3A,B**). Here we show that male pECD mice have more fragmented sleep during the dark period, with an increase in time spent sleeping in short bout bins (32 and 64s) and a trend toward less time spent in larger sleep bins (256s; **Fig. 3A,B**). There was no difference in total bouts during the light or dark period (**Fig. 3C,D**). However, male mice did display shorter bout lengths, specifically during the dark period (**Fig. 3B,E,F**). This reduction in dark bout length suggests more fragmented sleep. Along with this decrease in sleep bout length during the dark, male pECD mice showed a trend for more sleep during the light period, with no change in total sleep during the dark period (**Fig.3G,H**). Together these data show that pECD increases the number of short sleep bouts in male mice.

**Figure 3.**
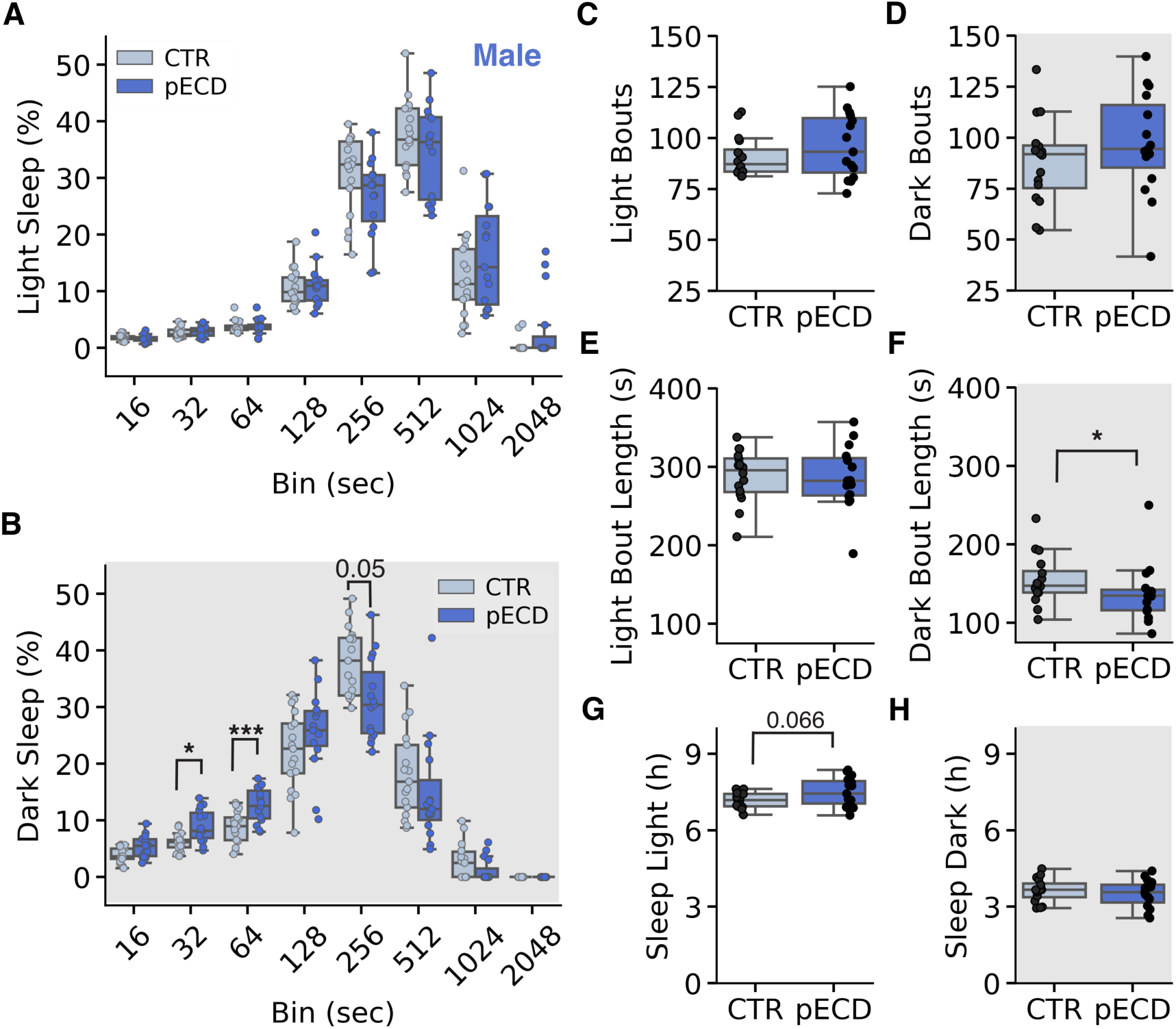
Male mice show a more fragmented sleep phenotype following pECD. ***A,*** Percent time spent asleep distributed into bout lengths during the light (Treatment: F_(1,30)_ = 4.33, *p* = 0.048; Bin: F_(7,210)_ = 123.02, *p* < 0.0001) and (***B***) dark (Treatment: F_(1,30)_ = 2.20, *p* = 0.151; Bin: F_(7,210)_ = 122.96, *p* < 0.0001) phases. ***C,*** Boxplot of total number of sleep bouts during the light (*inactive;* t(29) = 1.13, *p* = 0.27) and (***D***) dark (*active*; t(29) = 1.07, *p* = 0.29) period. ***E,*** Bout length (s) split between light (t(28) = 0.44, *p* = 0.66) and ***F,*** dark (t(27) = 2.28, *p* = 0.03). ***G,*** Total sleep time (h) during the light (t(29) = 1.91, *p* = 0.066) and (***H***) dark period (t(29) = 0.78, *p* = 0.44). Data are averages of 3 days for each mouse (*n* = 14-17 mice/group). *Two-way ANOVA and student t-test*.

### pECD does not impact sleep distribution or behavior in adult female pECD offspring

Previous work from our group and others have demonstrated that female mice are more resilient than male mice to certain changes in environmental factors, such as stress and time of day^6,18^. Similar to **Fig. 3**, we measured how female mice distributed their sleep within bins between the light and dark phases (**Fig. 4A,B**). In female mice we found no discernable differences between how mice distributed their sleep during the light and dark phases (**Fig. 4A,B**). These results were consistent when basic sleep measures were summarized into the light and dark phases. Specifically, there were no diurnal differences in total number of sleep bouts (**Fig. 4C,D**), average bout length (**Fig. 4E,F**), or total sleep time (**Fig. 4G,H**). Together, this suggests that pECD does not alter sleep structure in adult female mice.

**Figure 4.**
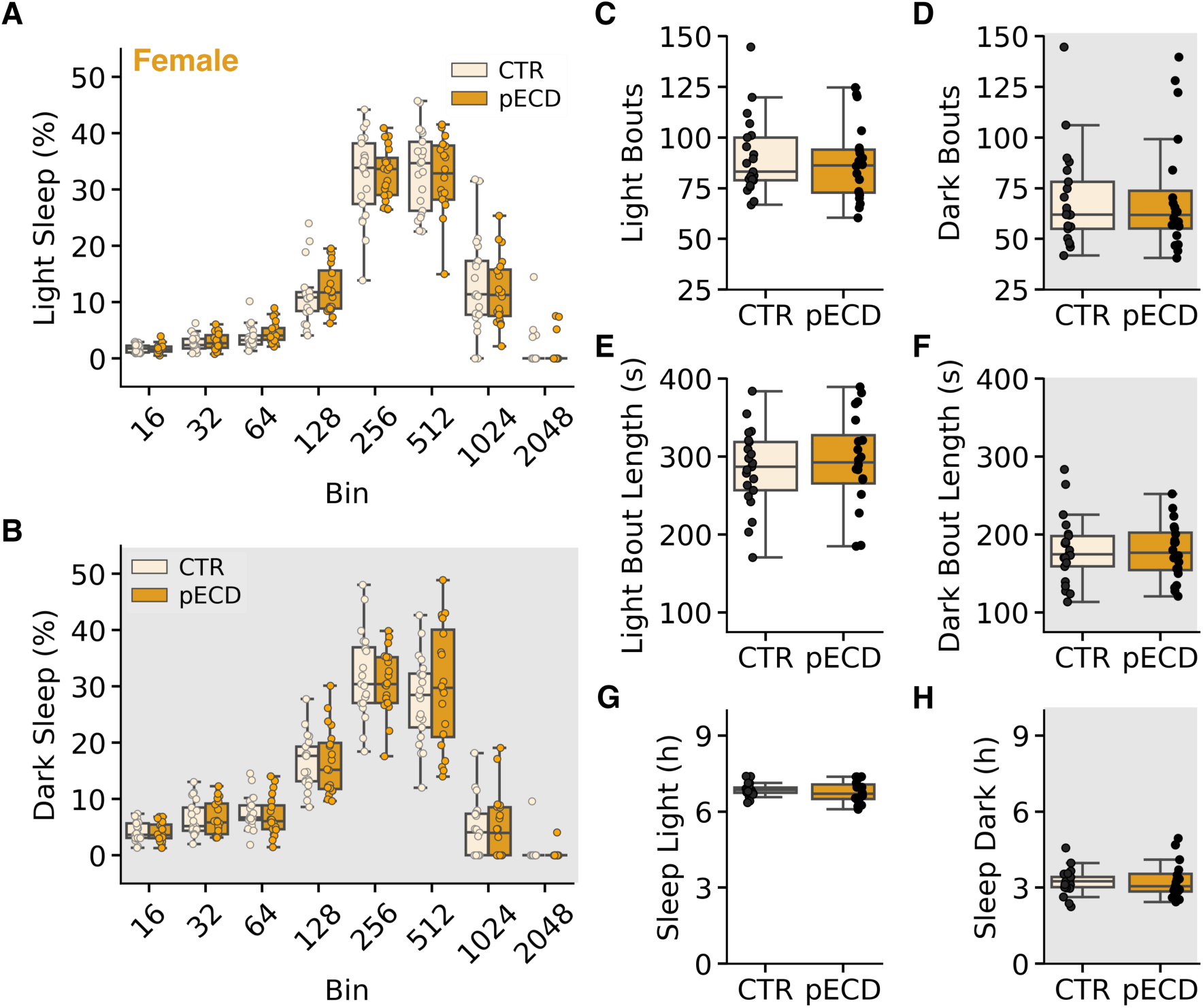
pECD does not impact sleep structure in females. ***A,*** Percent time spent asleep distributed into bout lengths during the light (Treatment: F_(1,35)_ = 0.29, *p* = 0.60; Bin: F_(7,245)_ = 266.77, *p* < 0.001) and (***B***) dark phases (Treatment: F_(1, 35)_ = 1.86, *p* = 0.18; Bin: F_(7, 245)_ = 171.75, *p* < 0.001). ***C,*** Boxplot of total number of sleep bouts during the light (*inactive;* t(38) = 0.03, *p* = 0.97) and (***D***) dark phases (*active*; t(35) = 0.87, *p* = 0.39). ***E,*** Bout length (s) split between light (t(39) = 0.44, *p* = 0.67) and ***F,*** dark (t(38) = 0.29, *p* = 0.78). ***G,*** Total sleep time (h) during the light (t(39) = 0.90, *p* = 0.37) and (***H***) dark phases (t(35) = 0.95, *p* = 0.35). Data are averages of 3 days for each mouse. (*n* = 20-21 mice/group). *Two-way ANOVA and Student t-test*.

### pECD alters postsynaptic component of excitatory inputs onto pyramidal PFC neurons in male and female mice

Sleep quality and neural function are tightly intertwined. The PFC is a region essential to cognitive function, learning and memory, and emotional regulation ^4,5,19,20^. Our previous work has demonstrated that basal properties of pyramidal neurons in layer 2/3 of the mPFC are rhythmic and ECD during adulthood drastically alters postsynaptic properties in these neurons independent of time of day^6^. Since pECD induced fractured sleep during the night and increased sleep during the light period, we wanted to determine how pECD impacts PFC function at the synaptic level. We measured excitatory postsynaptic currents (EPSCs) in female and male CTR and ECD mice at ZT 6-10 and 12-16 (**Fig. 5A,B**). We did not observe any effect of time of day or pECD on sEPSC frequency in either sex (**Fig. 5C,D**). However, there were numerous effects on postsynaptic components of synaptic transmission. In female CTR mice, there was increased amplitude and decreased decay time and half-width of sEPSCs at ZT 12-16, all of which were lost in pECD mice (**Fig. 5E,G,I**). These components contribute to the total sEPSC area, and given their inverse changes with time of day (higher amplitude, but shorter decay), there was no change in total sEPSC area (**Fig. 5K**). In male mice, there was no time of day effect on any postsynaptic components of sEPSCs, with the exception of total area (**Fig. 5F,H,J,L**). However, pECD had a large main effect on decay time, half-width, and total EPSC area (**Fig. 5H,J,L**). Combined, this data shows diurnal changes in the postsynaptic components of excitatory inputs in female mice, and pECD alters numerous postsynaptic components in both male and female mice. This strongly suggests that pECD impacts the function of these neurons, potential through changes in ion channel and receptor kinetics or expression.

**Figure 5.**
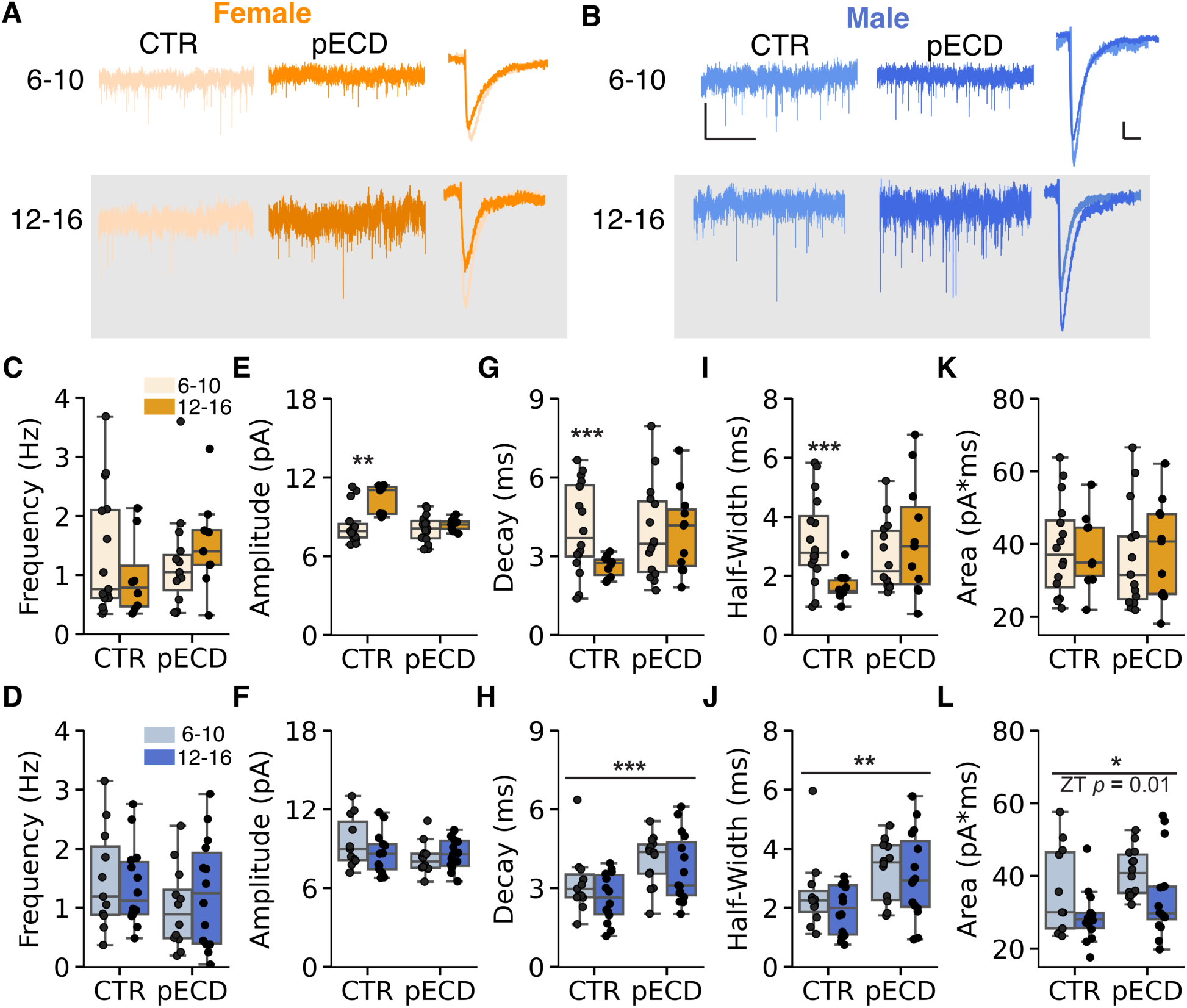
pECD and sex interact to alter mPFC excitatory inputs to pyramidal neurons. ***A*,** Representative traces of sEPSC voltage clamp recordings (*left*) and averaged event traces (*right*) overlayed from female and (***B***) male CTR and pECD mice at each ZT bin. ***C*,** Boxplot of sEPSC frequency in female (Treatment: F_(3, 43)_ = 0.113, *p* = 0.738; ZT: F_(3, 43)_ = 0.006, *p* = 0.938; Interaction: F_(3, 43)_ = 1.817, *p* = 0.185) and (***D***) male (Treatment: F_(3, 48)_ = 0.539, *p* = 0.466; ZT: F_(3, 48)_ = 0.024, *p* = 0.877; Interaction: F_(3, 48)_ = 0.103, *p* = 0.749) mice. ***E***, Boxplot of sEPSC amplitude in female (Treatment: F_(3, 44)_ = 6.927, *p* = 0.0117; ZT: F_(3, 44)_ = 9.874, *p* = 0.003; Interaction: F_(3, 44)_ = 6.376, *p* = 0.015) and (***F***) male (Treatment: F_(3, 46)_ = 1.118, p = 0.296; ZT: F_(3, 46)_ = 0.017, *p* = 0.898; Interaction: F_(3, 46)_ = 1.184, p = 0.282) mice at ZT6-10 and 12-16. ***G*,** Boxplot of sEPSC decay time in female (Treatment: F_(3, 48)_ = 1.078, *p* = 0.304; ZT: F_(3, 48)_ = 2.498, *p* = 0.121; Interaction: F_(3, 48)_ = 2.725, *p* = 0.105) and (***H***) male (Treatment: F_(3, 51)_ = 11.676, *p* = 0.0013; ZT: F_(3, 51)_ = 0.989, *p* = 0.325; Interaction: F_(3, 51)_ = 0.431, *p* = 0.515) mice at ZT6-10 and 12-16. ***I*,** Boxplot of sEPSC half-width in female (Treatment: F_(3, 47)_ = 0.850, *p* = 0.361; ZT: F_(3, 47)_ = 1.621, *p* = 0.209; Interaction: F_(3, 47)_ = 7.163, *p* = 0.010) and (***J***) male (Treatment: F_(3, 50)_ = 9.081, *p* = 0.004; ZT: F_(3, 50)_ = 1.067, *p* = 0.307; Interaction: F_(3, 50)_ = 0.259, *p* = 0.6129) mice at ZT6-10 and 12-16. ***K*,** Boxplot of sEPSC area in female (Treatment: F_(3, 48)_ = 0.199, *p* = 0.658; ZT: F_(3, 48)_ = 0.048, *p* = 0.827; Interaction: F_(3, 48)_ = 0.214, *p* = 0.646) and (***L***) male (Treatment: F_(3, 51)_ = 4.062, *p* = 0.049; ZT: F_(3, 51)_ = 7.317, *p* = 0.009; Interaction: F_(3, 51)_ = 0.003, *p* = 0.957) mice at ZT6-10 and 12-16 (*n* = 14-22 cells/group). Box plots represent median, min/max, and second and third quartiles. Two-way ANOVA with Tukey post-hoc analysis for Treatment and ZT bin, \**p* < 0.05, \*\**p* < 0.01, \*\*\**p* < 0.001.

### pECD does not alter action potential dynamics of pyramidal PFC neurons in female mice

Our results demonstrate that the effects of pECD on pyramidal neuron function in female mice are likely postsynaptic (**Fig. 5**). Our previous work shows that ECD in adulthood alters numerous action potential properties in male mice^6^. To better understand how pECD impacts cell endogenous traits we measured basal membrane properties and used a current-step protocol to evoke action potentials in these neurons (**Fig. 6A,B**). We found that pECD increased the membrane resistance (Rm) in these neurons but had no effect on membrane capacitance (Cm) or resting membrane potential (RMP) (**Fig. 6C-E**). Here we show that in female mice, pECD has no effect on the threshold for action potential firing or after hyperpolarization (AHP), but there was a trend in the interaction between treatment and ZT time action potential amplitude, with amplitude trending to increase during the dark period in CTR and decrease in pECD mice (**Fig. 6F-H**). We did not observe any pECD on decay time, action potential half-width, or area (**Fig. 6I-K**). Combined, this data suggests that pECD has little impact on cell endogenous traits and action potential dynamics in PFC pyramidal neurons of female mice.

**Figure 6.**
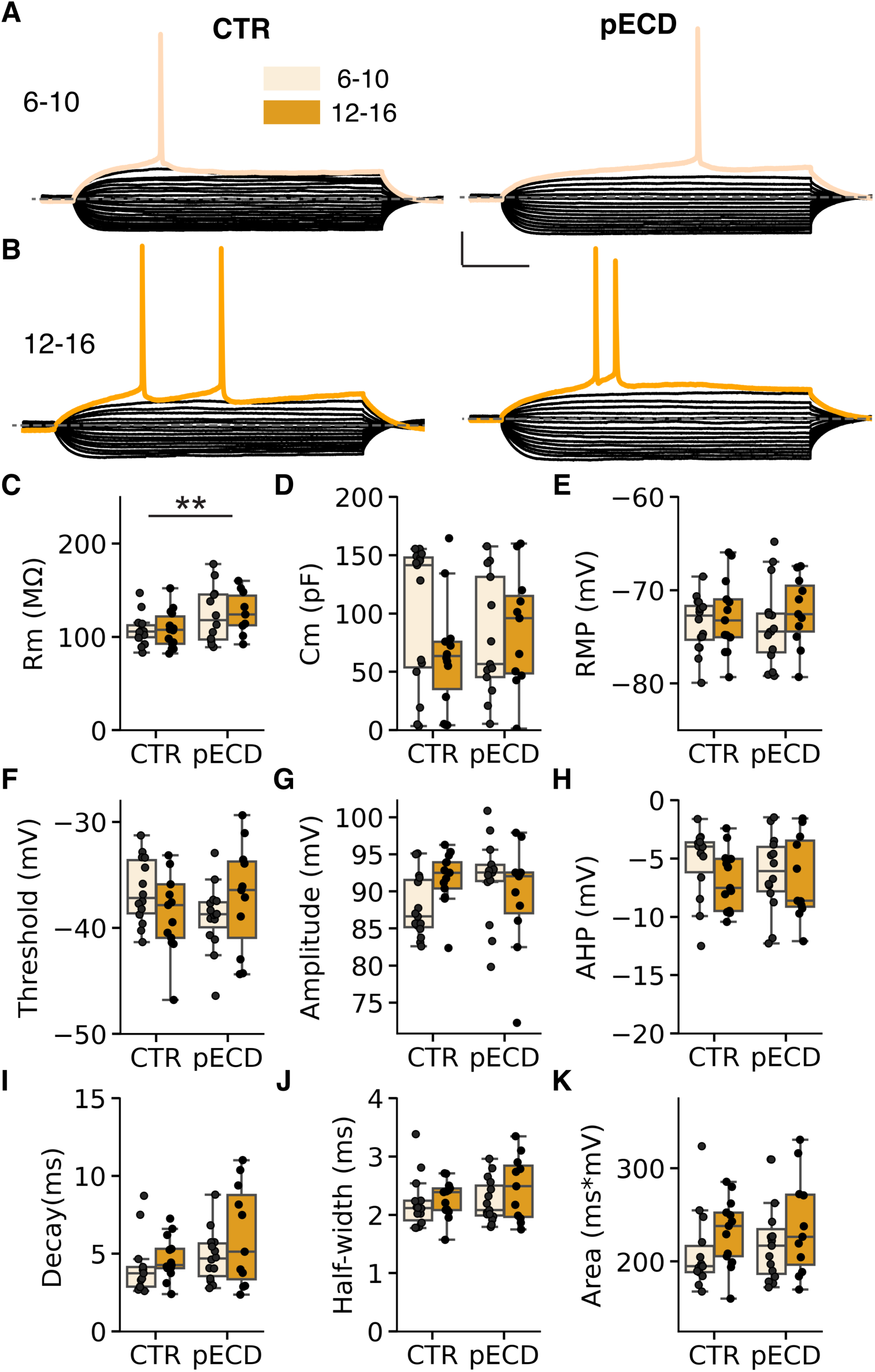
pECD alters cell endogenous properties of pyramidal PFC neurons in female mice. ***A,*** Representative trace of evoked action potentials in mPFC pyramidal neurons of female CTR and (***B***) pECD mice. ***C,*** Boxplot of membrane resistance (Treatment: F_(3, 46)_ = 7.379, *p* = 0.009; ZT: F_(3, 46)_ = 0.0449, *p* = 0.833; Interaction: F_(3, 46)_ = 0.006, *p* = 0.940), ***D*,** capacitance (pF) (Treatment: F_(3, 49)_ = 0.002, *p* = 0.967; ZT: F_(3, 49)_ = 1.425, *p* = 0.238; Interaction: F_(3, 49)_ = 2.404, *p* = 0.128), ***E,*** resting membrane potential (RMP) (Treatment: F_(3, 51)_ = 0.02, *p* = 0.888; ZT: F_(3, 51)_ = 1.105, *p* = 0.298; Interaction: F_(3, 51)_ = 0.088, *p* = 0.768), ***F,*** action potential (AP) threshold (Treatment: F_(3, 49)_ = 0.792, *p* = 0.378; ZT: F_(3, 49)_ = 0.034, *p* = 0.854; Interaction: F_(3, 49)_ = 2.029, *p* = 0.161), ***G,*** amplitude (Treatment: F_(3, 49)_ = 2.376, *p* = 0.130; ZT: F_(3, 49)_ = 2.139, *p* = 0.15; Interaction: F_(3, 49)_ = 3.269, *p* = 0.077), ***H,*** afterhyperpolarization (AHP; Treatment: F_(3, 48)_ = 0.243, *p* = 0.624; ZT: F_(3, 48)_ = 1.435, *p* = 0.237; Interaction: F_(3, 48)_ = 0.420, *p* = 0.520), ***I,*** decay tau (Treatment: F_(3, 47)_ = 3.037, *p* = 0.088; ZT: F_(3, 47)_ = 2.306, *p* = 0.136; Interaction: F_(3, 47)_ = 0.470, *p* = 0.496), ***J,*** half-width (ms) (Treatment: F_(3, 49)_ = 0.824, *p* = 0.369; ZT: F_(3, 49)_ = 1.541, *p* = 0.221; Interaction: F_(3, 49)_ = 0.427, *p* = 0.516), and ***K,*** total area (ms*mV) (Treatment: F_(3, 49)_ = 0.614, *p* = 0.437; ZT: F_(3, 49)_ = 2.552, *p* = 0.117; Interaction: F_(3, 49)_ = 0.037, *p* = 0.849) from CTR and pECD female mice at ZT 6-10 and 12-16 (*n* = 14-22 cells/group). *Two-way ANOVA for main effects of treatment, time, and interactions.* \*\**p* < 0.01.

### pECD alters cell endogenous properties of pyramidal PFC neurons in male mice

Similar to our synaptic findings in female mice, male mice also displayed the largest pECD effects on the postsynaptic components of excitatory inputs, such as decay time, half-width, and EPSC area (**Fig. 5H-L**). To probe if pECD alters the cell endogenous traits of plPFC pyramidal neurons we measured basal properties and used the same current step protocol as in **Fig. 6** to measure action potential dynamics (**Fig. 7A-B, F-K**). There was no impact of time-of-day or pECD on membrane resistance, but pECD induced a loss of diurnal changes in both membrane capacitance and RMP (**Fig. 7C-E**). Similarly, pECD did not have a significant impact on action potential dynamics (**Fig. 7F-K**). However, there were trends to a strengthened diurnal difference in action potential threshold (**Fig. 7F, J-K**). In combination with our sleep experiments (**Fig. 3D, F, G**), these results suggest that pECD has the largest impact on behavior and neural function during the dark period.

**Figure 7.**
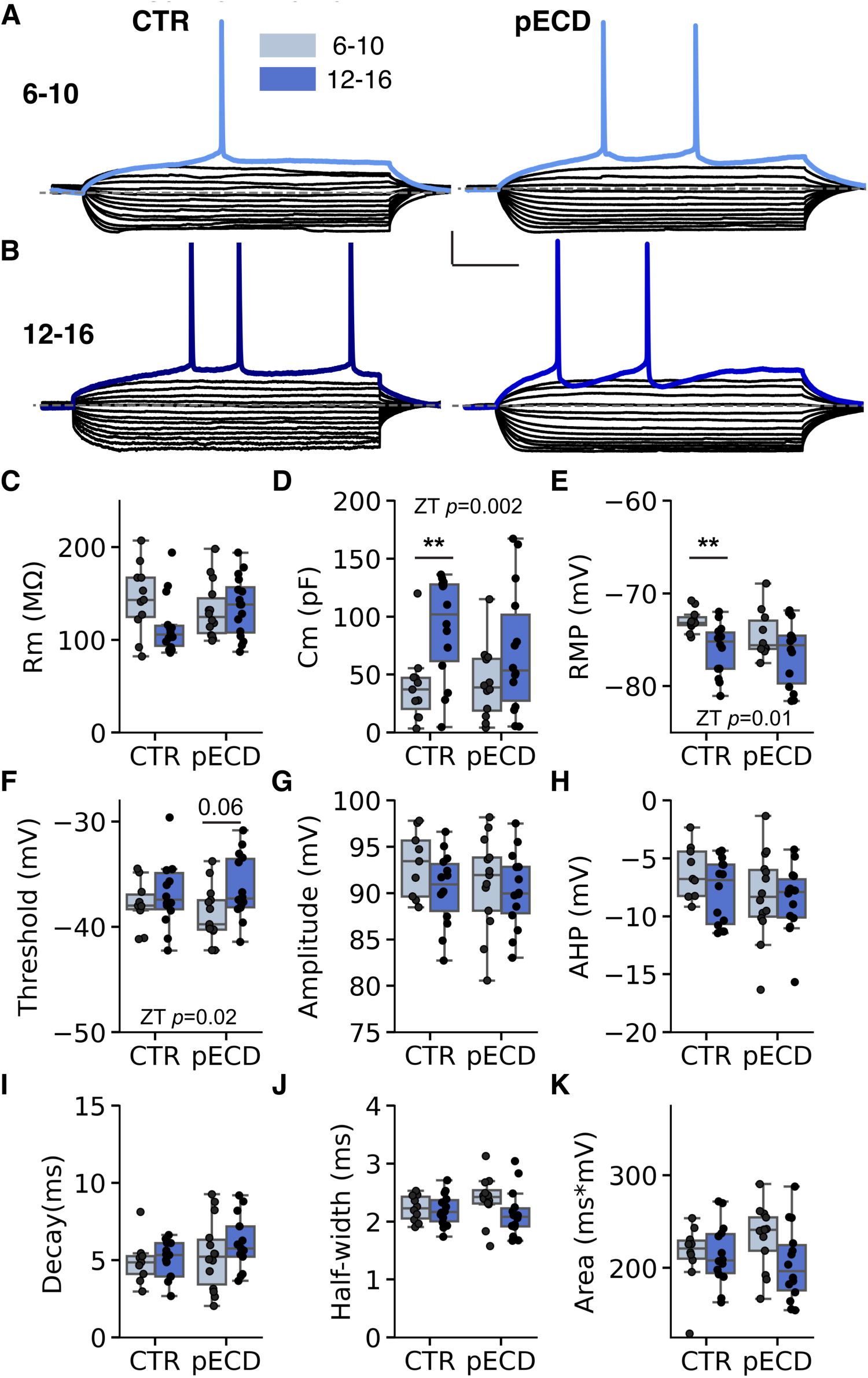
pECD alters cell endogenous properties of pyramidal PFC neurons in male mice. ***A,*** Representative trace of evoked action potentials in mPFC pyramidal neurons of male CTR and (***B***) pECD mice. ***C,*** Boxplot of membrane resistance (Treatment: F_(3, 51)_ = 0.4111, *p* = 0.5243; ZT: F_(3, 51)_ = 1.4672, *p* = 0.2314; Interaction: F_(3, 47)_ = 3.560, *p* = 0.065), ***D*,** capacitance (pF) (Treatment: F_(3, 51)_ = 0.705, *p* = 0.405; ZT: F_(3, 51)_ = 10.544, *p* = 0.002; Interaction: F_(3, 51)_ = 1.016, *p* = 0.319), ***E,*** resting membrane potential (RMP) (Treatment: F_(3, 46)_ = 2.17, *p* = 0.148; ZT: F_(3, 46)_ = 7.108, *p* = 0.011; Interaction: F_(3, 46)_ = 1.045, *p* = 0.312), ***F,*** action potential (AP) threshold (Treatment: F_(3, 48)_ = 0.146, *p* = 0.704; ZT: F_(3, 48)_ = 6.154, *p* = 0.017; Interaction: F_(3, 48)_ = 1.557, *p* = 0.218), ***G,*** amplitude (Treatment: F_(3, 46)_ = 0.696, *p* = 0.409; ZT: F_(3, 46)_ = 1.816, *p* = 0.185; Interaction: F_(3, 46)_ = 0.567, *p* = 0.455), ***H,*** afterhyperpolarization (Treatment: F_(3, 47)_ = 2.208, *p* = 0.144; ZT: F_(3, 47)_ = 1.216, *p* = 0.276; Interaction: F_(3, 47)_ = 0.541, *p* = 0.466), ***I,*** decay tau (Treatment: F_(3, 45)_ = 2.250, *p* = 0.141; ZT: F_(3, 45)_ = 0.943, *p* = 0.337; Interaction: F_(3, 45)_ = 0.407, *p* = 0.527), ***J,*** half-width (ms) (Treatment: F_(3, 46)_ = 0.151, *p* = 0.700; ZT: F_(3, 46)_ = 2.322, *p* = 0.134; Interaction: F_(3, 46)_ = 1.309, *p* = 0.258), and ***K,*** total area (ms*mV) (Treatment: F_(3, 47)_ = 0.078, *p* = 0.781; ZT: F_(3, 47)_ = 2.222, *p* = 0.143; Interaction: F_(3, 47)_ = 2.00, *p* = 0.164) CTR and pECD male mice at ZT 6-10 and 12-16 (*n* = 11-22 cells/group). *Two-way ANOVA for main effects of treatment, time, and interactions.* \*\**p* < 0.01.

## Discussion

Here we demonstrate three main findings. First, pECD decreases offspring body weight, an effect that is sustained into adulthood but only in male mice. Second, pECD causes fragmented sleep in males while trending toward increased sleep in the light period. Lastly, we show that pECD alters synaptic and cell endogenous function of PFC pyramidal neurons in both male and female mice. This pattern of responses indicates that the impact of pECD is sexually dimorphic, and that the impacts on sleep can be distinct from the impact on synaptic function.

Metabolic function and daily rhythms are highly intertwined^4,21,22^. Adult mice subjected to circadian disruption steadily gain body weight over time^4^. However, pECD seems to lead to a blunting of weight gain early in life, an effect that is sustained in males, but seems to dissipate in females. One potential explanation is that there are multiple mechanisms underlying metabolic regulation, some of which are both underdeveloped and have inverse actions early in life^23–26^. For example, hypothalamic signaling of the satiety hormone leptin does not fully develop until the second to third week of life^23,24,27^. Thus, these developmental differences in metabolic regulation may underly why pECD results in a decrease in body weight. Alternatively, it is possible that pECD decreases the gestational period and these pups are born prematurely. While we did not observe differences in litter size, nor a difference at P21, future studies using timed pregnancy are necessary to better probe this potential outcome. This would allow for a disentangling of the metabolic components of pECD, and potential functional changes are occurring during development during and after the gestational period.

Sleep behavior is complex, and mice are polyphasic sleepers, sleeping in short bouts on the order of minutes. We have shown that ECD in adults leads to a decrease in sleep quality (as measured by delta power) during the period of T20 environmental desynchronization^28^. Thus, while T20 ECD in adults does not lead to sleep loss, it does seem to reduce sleep quality in male mice^28^. Although in the present study we observed no effects on the total time mice spent asleep during the 24h period following pECD, as noted above, amount of sleep is not synonymous with sleep quality. There was a clear decrease in sleep bout length during the dark period in male mice. This suggests that pECD may lead to more fragmented sleep later in life, which similar to sleep disruption, is associated with poor health outcomes^17,29^. However, the dark period is when mice are most active. We speculate two potential outcomes of pECD leading to these outcomes. First, it is possible that male pECD mice are getting more restful sleep during the inactive period, resulting is less need for longer “naps” during the active period. Alternatively, pECD mice may sleep more during the inactive period because they are not receiving the normal rest and/or siesta common to control mice during the active period. While our non-invasive sleep measurement approach has many benefits, to assess the mechanisms that drive these changes in sleep structure, as well as sleep quality, we would need to apply more invasive EEG approaches.

Recent work from our lab shows that physiological properties of PFC pyramidal neurons changes throughout the 24hr day and ECD alters the fundamental function and morphology of these neurons^4,6^. In our present study, female CTR mice showed diurnal changes in postsynaptic EPSC components. Specifically, we saw an increase in amplitude, which may indicate increases in the number of glutamate receptors that can be activated. Additionally, the kinetics of these receptors has likely changed, as a decrease in half-width and decay time indicates that these channels are opening and closer much faster during the active period. Although, we observed no effect of pECD on sleep in female mice, we did note that time of day effects on EPSC characteristics were not present in pECD mice, which suggests that independent of sleep, pECD affects neural physiology and may impact other behaviors related to cognitive function or learning and memory. In male mice, pECD increased the overall area of EPSCs, due to increases in half-width and decay time. Opposite of female mice, this suggests that pECD alters the overall kinetics of glutamatergic signaling and these receptors may stay activated for longer in male ECD mice.

ECD in adult mice seems to abolish the impact of time-of-day and alters nearly all components of action potential dynamics^6^. Here we show that pECD also blunts diurnal changes in cell endogenous function. For example, we show that male pECD mice did not show diurnal changes in membrane capacitance or RMP as observed in CTR mice. Membrane capacitance is often associated with cell size, which is in part regulated by local ion concentrations that alter osmolarity and can influence the RMP. Local ion concentrations of Na+, K+, Ca2+ in the cortex, and cytosolic Ca2+ in the SCN, change with time of day^30,31^. This is one potential mechanism that could explain both our synaptic and cell endogenous findings. However, future studies investigating how pECD blunts diurnal changes in cell endogenous function are necessary to fully understand the broader implications of this finding. Although there was a trend toward diurnal changes in action potential threshold in pECD mice, this is not surprising as there was a strong main effect of time-of-day, similar to our previous work^1^. Female mice did not display diurnal changes in resting properties or action potential dynamics and similar to our sleep data, pECD had little impact on these measures. Of note, female pECD mice had a rebound in body weight late in development. This raises the question of how gonadal (or other) hormones during puberty may influence sleep and neural function in adulthood. An important consideration for future work is whether pECD may alter sleep and neural function during early development in female mice and diverge from their male counterparts after the onset of puberty.

Together, we conclude that pECD has long-lasting consequences for body weight, sleep, and mPFC synaptic function. Future studies can now focus on mechanisms by which disrupted day-night cycles can result in sustained changes even long after mice have been returned to normal environmental conditions. Although the effects of pECD were not as striking as those found with ECD in adulthood, our work highlights how perturbations in the developmental period can have lifelong impacts on brain and behavior. Further, an important area of investigation is to determine how pECD impacts physiological responses to additive environmental perturbations, such as susceptibility to a high-fat diet, stress responses, and circadian disruption later in life. This work forms an important foundation for these future studies and underscores that environment during the developmental period has numerous lifelong impacts on cellular and whole-organismal physiology.

## Methods

### Animals

All experiments and animal procedures were approved by the University of Massachusetts Amherst Institutional Care and Use Committee in accordance with the U.S. Public Health Service Policy on Humane Care and Use of Laboratory Animals and the National Institutes of Health *Guide for the Care and Use of Laboratory Animals* and are in compliance with ARRIVE guidelines^32^. Male and female NPY-GFP mice (Strain #006417, The Jackson Laboratory, Bar Harbor, ME, USA) on a C57BL/6J background were bred in-house used for these studies. All mice were group-housed in light-tight housing boxes at 25°C, under a 12:12-hr light:dark (LD; T24) cycle, with food and water available *ad libitum*. LD cycles in housing boxes were offset so that experiments from each zeitgeist time (ZT) bin occurred at the same external (real world) time each day. For electrophysiology studies mice were anesthetized in a chamber with isoflurane before euthanasia by decapitation.

### Maternal circadian desynchronization

For maternal circadian desynchronization experiments, dams (8-10 weeks of age) were placed in a 10:10 LD cycle (T20) for two weeks prior to being paired for breeding. Male breeders were maintained in a standard 12:12 LD (T24) cycle and placed with the female for 3-5 days before being placed back into standard T24 conditions. To control for multiparity, only second and third litters were used from each dam. Dams remained in T24 conditions for their first litter, at which point pups were removed and dams were placed into the above ECD conditions for the remainder of the experiment. To control for metabolic impacts on litter size (i.e. over and under nutrition) only litters with 6-9 pups were used. Litters with >9 pups were culled to 9 pups on P3. At postnatal day 21, pups were weaned into group housed cages with 2-3 same sex littermates per cage. Cages were randomly assigned to a lightbox ZT time and body weights were measured by hand every 4-7 days until P47 (adulthood).

### Sleep recordings

Sleep behavior was measured using a non-invasive piezo-sleep system (Signal Solutions LLC, Lexington, KY, USA). Mice were single housed in the sleep system for 5-6 days with a piezo apparatus under the cage floor, had *ad libitum* access to food and water and were left undisturbed for the recording period. The first 48h were considered as a habituation period and only the last 72h of sleep recordings were used for analysis. Data was acquired using the PiezoSleep software. Sleep behavior was calculated in SleepStats 2p151 (Signal Solutions LLC, Lexington, KY, USA) and exported as a .csv file for further statistical analysis. After sleep recordings, mice remained in single-housed conditions until utilized for electrophysiology experiments.

### Electrophysiology

Two mice were simultaneously euthanized 1h prior to their ZT bin (i.e., mice were euthanized at ZT11 for recording bin ZT12-16). Due to the timing of our recording bins no mice were sacrificed during the dark period. After euthanasia, brains were immediately removed and the forebrain was blocked while bathing in a 0-4°C oxygenated N-methyl-D-glucamine (NMDG) - 4-(2-hydroxyethyl)-1-piperazineethanesulfonic acid (HEPES) cutting solution composed of (mM): 92 NMDG, 2.5 KCl, 1.25 NaH_2_PO_4_, 30 NaHCO_3_, 3 sodium pyruvate, 2 thiourea, 20 HEPES, 10 MgSO_4_, 0.5 CaCl_2_, 25 glucose, 20 sucrose. Cutting solution was brought to pH 7.4 with ∼17mL of 5M HCl ^33^. The forebrains were mounted adjacent to each other and sectioned simultaneously on a vibratome (VT1200S, Leica Biosciences, Buffalo Grove, IL, USA) with a sapphire knife (Delaware Diamond Knives, Wilmington, DE, USA) yielding roughly three slices containing the PFC from each (250-μm) per mouse. Slices were transferred and allowed to recover for 30-45 min in room temperature recording artificial cerebrospinal fluid (aCSF) solution composed of (mM): 124 NaCl, 3.7 KCl, 2.6 NaH_2_PO_4_, 26 NaHCO_3_, 2 CaCl_2_, 2 MgSO_4_, 10 glucose. aCSF had a final pH of 7.3-7.4, osmolarity of 307-310 mOsmos, and was continuously bubbled using 95% 0_2_/5% C0_2_. For recordings, brain slices were transferred to a perfusion chamber containing aCSF maintained at 34-37°C with a flow rate of 1mL/min. Neurons were visualized using an upright microscope (Zeiss Axoskop 2, Oberkochen, Germany). Recording electrodes were back-filled with experiment-specific internal solutions as follows (mM): Current-clamp and voltage-clamp; 125 K-gluconate, 10 KCl, 10 NaCl, 5 HEPES, 10 EGTA, 1 MgCl_2_, 3 NaATP and 0.25 NaGTP (liquid-junction potential (LJP) = ∼14.5 mV; Predicted E_K_ = ∼-95 mV). All internal solutions were brought to pH 7.3 using KOH at 301-304 mOsm. Patch electrodes with a resistance of 4-6MΩ were guided to neurons with an MPC-200-ROE controller and MP285 mechanical manipulator (Sutter Instruments, Novato, CA, USA). Patch-clamp recordings were collected through a UPC-10 USB dual digital amplifier and Patchmaster NEXT recording software (HEKA Elektronik GmbH, Reutlingen, Germany). Current clamp voltage-step protocols were performed from a starting V_H_ = -70 mV and used 1s 10-pA steps from -80 pA to +230 pA. All compounds were obtained from Tocris Cookson, Cayman Chemical, and Sigma Aldrich.

A small percentage (∼10%) of all recorded neurons had unique characteristics in resting membrane properties and action potential dynamics that were independent of time-of-day, and analysis was limited to our previously defined and characterized Type I pyramidal neurons^6^.

### Statistical Analysis

Only neurons with input resistance > 70 MΩ were studied. Neurons were not considered for further analysis if series resistance exceeded 50MΩ or drifted >10% during baseline. Rheobase was calculated as the first current step to elicit an action potential and action potential dynamics (threshold, decay tau, and half-width) were obtained from the first evoked action potential to avoid variance in ion channel function due to repeated action potential firing. G*Power 3.0 software (Franz Faul, Uni Kiel, Germany) was used to conduct our power analysis, for a *p* value of <0.05 with 90% power. Adequate sample sizes were based upon expected effect sizes from similar experiments. Raw data files were analyzed in the Patchmaster NEXT software or converted using ABF Utility (Synaptosoft) for analysis in MiniAnalysis (Synaptosoft). N-values for analysis and presented in figures represent individual cells. Where applicable, extreme outliers were identified using the ROUT method with a conservative Q = 0.5%. To control for biological variability between groups N = 4-8 mice per group was used. To control for within animal variability 2-3 brain slices were collected per mouse. For experiments including the use of drug, only one cell per slice was used. Statistical comparison of effects between each time-period was made using a full model two-way ANOVA (column, row, and interaction effects) unless otherwise noted. All calculations and statistical analyses were performed in Python 3.11.3 using the JupyterLab 3.6.3 ^34^. NumPy ^35^ and Pandas^36^ packages were used for data handling, computations, and organization. Statistics were performed with Pingouin 0.5.3 ^37^ package and data visualization was created using Matplotlib 3.7.1 ^38^ and Seaborn 0.12 ^39^ packages.

## References

1. Lauby, S. C., Fleming, A. S. & McGowan, P. O. Beyond maternal care: The effects of extra-maternal influences within the maternal environment on offspring neurodevelopment and later-life behavior. Neurosci. Biobehav. Rev. 127, 492–501 (2021).

2. Panera, N. et al. Genetics, epigenetics and transgenerational transmission of obesity in children. Front. Endocrinol. 13, 1006008 (2022).

3. Amrom, D. & Schwartz, S. S. Maternal Metabolic Health, Lifestyle, and Environment - Understanding How Epigenetics Drives Future Offspring Health. Curr. Diabetes Rev. 19, e220422203919 (2023).

4. Karatsoreos, I. N., Bhagat, S., Bloss, E. B., Morrison, J. H. & McEwen, B. S. Disruption of circadian clocks has ramifications for metabolism, brain, and behavior. Proc. Natl. Acad. Sci. 108, 1657–1662 (2011).

5. Miller, E. K. & Cohen, J. D. An Integrative Theory of Prefrontal Cortex Function. Annu. Rev. Neurosci. 24, 167–202 (2001).

6. Roberts, B. L. & Karatsoreos, I. N. Circadian desynchronization disrupts physiological rhythms of prefrontal cortex pyramidal neurons in mice. Sci. Rep. 13, 9181 (2023).

7. Lorsung, E., Karthikeyan, R. & Cao, R. Biological Timing and Neurodevelopmental Disorders: A Role for Circadian Dysfunction in Autism Spectrum Disorders. Front. Neurosci. 15, 642745 (2021).

8. Wallace, N. K., Pollard, F., Savenkova, M. & Karatsoreos, I. N. Effect of Aging on Daily Rhythms of Lactate Metabolism in the Medial Prefrontal Cortex of Male Mice. Neuroscience (2020) doi:10.1016/j.neuroscience.2020.07.032.

9. McEwen, B. S. & Morrison, J. H. Brain On Stress: Vulnerability and Plasticity of the Prefrontal Cortex Over the Life Course. Neuron 79, 16–29 (2013).

10. Irwin, M. R. Why sleep is important for health: a psychoneuroimmunology perspective. Annu. Rev. Psychol. 66, 143–172 (2015).

11. Yuen, E. Y. et al. Acute stress enhances glutamatergic transmission in prefrontal cortex and facilitates working memory. Proc. Natl. Acad. Sci. 106, 14075–14079 (2009).

12. Moorman, D. E., James, M. H., McGlinchey, E. M. & Aston-Jones, G. Differential roles of medial prefrontal subregions in the regulation of drug seeking. Brain Res. 1628, 130–146 (2015).

13. Zaitsev, A. V., Povysheva, N. V., Gonzalez-Burgos, G. & Lewis, D. A. Electrophysiological classes of layer 2/3 pyramidal cells in monkey prefrontal cortex. J. Neurophysiol. 108, 595–609 (2012).

14. Radnikow, G. & Feldmeyer, D. Layer- and Cell Type-Specific Modulation of Excitatory Neuronal Activity in the Neocortex. Front. Neuroanat. 12, (2018).

15. Topchiy, I., Fink, A. M., Maki, K. A. & Calik, M. W. Validation of PiezoSleep Scoring Against EEG/EMG Sleep Scoring in Rats. Nat. Sci. Sleep 14, 1877–1886 (2022).

16. Mang, G. M. et al. Evaluation of a Piezoelectric System as an Alternative to Electroencephalogram/ Electromyogram Recordings in Mouse Sleep Studies. Sleep 37, 1383–1392 (2014).

17. Wallace, E. et al. Differential effects of duration of sleep fragmentation on spatial learning and synaptic plasticity in pubertal mice. Brain Res. 1615, 116–128 (2015).

18. Yuen, E. Y., Wei, J. & Yan, Z. Estrogen in prefrontal cortex blocks stress-induced cognitive impairments in female rats. J. Steroid Biochem. Mol. Biol. 160, 221–226 (2016).

19. Woodruff, E. R. et al. Coordination between Prefrontal Cortex Clock Gene Expression and Corticosterone Contributes to Enhanced Conditioned Fear Extinction Recall. eNeuro 5, (2018).

20. Popoli, M., Yan, Z., McEwen, B. S. & Sanacora, G. The stressed synapse: the impact of stress and glucocorticoids on glutamate transmission. Nat. Rev. Neurosci. 13, 22–37 (2012).

21. Kohsaka, A. et al. High-Fat Diet Disrupts Behavioral and Molecular Circadian Rhythms in Mice. Cell Metab. 6, 414–421 (2007).

22. Padilla, S. L. et al. Kisspeptin Neurons in the Arcuate Nucleus of the Hypothalamus Orchestrate Circadian Rhythms and Metabolism. Curr. Biol. 29, 592–604.e4 (2019).

23. Roberts, B. L., Bennett, C. M., Carroll, J. M., Lindsley, S. R. & Kievit, P. Early overnutrition alters synaptic signaling and induces leptin resistance in arcuate proopiomelanocortin neurons. Physiol. Behav. 206, 166– 174 (2019).

24. Baquero, A. F. et al. Developmental switch of leptin signaling in arcuate nucleus neurons. J. Neurosci. Off. J. Soc. Neurosci. 34, 9982–9994 (2014).

25. Bouret, S. G., Draper, S. J. & Simerly, R. B. Formation of projection pathways from the arcuate nucleus of the hypothalamus to hypothalamic regions implicated in the neural control of feeding behavior in mice. J. Neurosci. Off. J. Soc. Neurosci. 24, 2797–2805 (2004).

26. Bouret, S. G., Bates, S. H., Chen, S., Myers, M. G. & Simerly, R. B. Distinct Roles for Specific Leptin Receptor Signals in the Development of Hypothalamic Feeding Circuits. J. Neurosci. 32, 1244–1252 (2012).

27. Glavas, M. M. et al. Early Overnutrition Results in Early-Onset Arcuate Leptin Resistance and Increased Sensitivity to High-Fat Diet. Endocrinology 151, 1598–1610 (2010).

28. Phillips, D. J., Savenkova, M. I. & Karatsoreos, I. N. Environmental disruption of the circadian clock leads to altered sleep and immune responses in mouse. Brain. Behav. Immun. 47, 14–23 (2015).

29. Xiong, Y. et al. Sleep fragmentation reduces explorative behaviors and impairs motor coordination in male mice. J. Neurosci. Res. 102, e25268 (2024).

30. Ikeda, M. et al. Circadian dynamics of cytosolic and nuclear Ca2+ in single suprachiasmatic nucleus neurons. Neuron 38, 253–263 (2003).

31. Bijak, M. Daily and seasonal variations in Na+, K+, Ca2+ and Mg2+ contents in the cingulate cortex of the mouse brain. Folia Biol. (Praha) 37, 3–11 (1989).

32. Sert, N. P. du et al. The ARRIVE guidelines 2.0: Updated guidelines for reporting animal research. PLOS Biol. 18, e3000410 (2020).

33. Ting, J. T. et al. Preparation of Acute Brain Slices Using an Optimized N-Methyl-D-glucamine Protective Recovery Method. JoVE J. Vis. Exp. e53825 (2018) doi:10.3791/53825.

34. GitHub - jupyterlab/jupyterlab: JupyterLab computational environment. https://github.com/jupyterlab/jupyterlab.

35. Harris, C. R. et al. Array programming with NumPy. Nature 585, 357–362 (2020).

36. team, T. pandas development. pandas-dev/pandas: Pandas. Zenodo 10.5281/zenodo.7857418 (2023).

37. Vallat, R. Pingouin: statistics in Python. J. Open Source Softw. 3, 1026 (2018).

38. Caswell, T. A., et al. matplotlib/matplotlib: REL: v3.7.1. Zenodo 10.5281/zenodo.7697899 (2023).

39. Waskom, M. L. seaborn: statistical data visualization. J. Open Source Softw. 6, 3021 (2021).

